# Integrating Longitudinal Metabolite Profiles Improves Trait Prediction in Pigs in a Trait- and Timepoint-Dependent Manner

**DOI:** 10.1101/2025.11.06.687040

**Authors:** Quazi Abir Hassan Roddur, Jiyai Qu, Dingzhen Liang, Eula Regina Carrara, Daniela Lourenco, Bruno Valente, Ching-Yi Chen, Justin Holl, Hao Cheng

## Abstract

**Background:** Accurate prediction of genetic merit is essential for accelerating genetic improvement in pigs, particularly for traits that are costly or difficult to measure directly. This study investigated the potential of integrating individual-level blood serum metabolite profiles sampled at two developmental stages (10-week and 20-week) into genomic prediction models for five economically important traits: average daily feed intake (DFI), feed conversion ratio (FCR), backfat thickness (BF), test daily gain (TDG), and loin depth (LD). Using a BayesC modeling framework, we analyzed 1,637 pigs from a single purebred population with complete phenotype, genotype, and metabolite-profile data. We evaluated seven models, including a genotype-only model (G), metabolite-only models using metabolite-profile data from 10-week (M1), 20-week (M2), or both timepoints (M1+M2), and combined models integrating genotypes with metabolite-profile data (G+M1, G+M2, G+M1+M2).

**Results:** Heritability estimates were generally low to moderate, ranging from 0.07 to 0.30 for the 10-week metabolite profile and from 0.04 to 0.28 for the 20-week metabolite profile. Prediction accuracy of phenotype consistently improved when metabolite-profile data were integrated into genotype-based models, although the magnitude of improvement varied depending on the trait and the timepoint of metabolite sampling. Prediction accuracy increased from 0.31 (G) to 0.41 (G+M2; G+M1+M2) for DFI, and from 0.27 (G) to 0.33 (G+M2; G+M1+M2) for FCR. The latter two models also delivered the largest gains over G for BF (from 0.41 to 0.45) and TDG (from 0.28 to 0.32). However, LD benefited the most when both 10-week and 20-week metabolite profiles were combined (G+M1+M2: 0.45 compared to G: 0.42).

**Conclusions:** Across all traits, models combining genotype data with metabolite profiles from one or multiple timepoints achieved the highest or equally high prediction accuracies compared to the genotype-only model, reflecting complementary biological insights captured by metabolite profiles. These findings highlight the potential value of metabolite-profile data as an intermediate omics layer to enhance genomic prediction, particularly when integration strategies are tailored to trait-specific biology and sampling timepoints.

## Background

Accurate prediction of genetic merits is essential for the genetic improvement of livestock (Bijma, 2012; Van der Werf, 2010). Genomic prediction approaches that leverage genotypes and phenotypes from relatives have become standard practice in animal breeding (Mrode et al., 2019; Wiggans et al., 2011) since Meuwissen et al. (2001) proposed genomic selection using simulated data. In pig breeding, traits of economic importance, such as feed efficiency, growth, and reproduction, are often expensive, labor-intensive, or challenging to measure directly in selection candidates (Guan et al., 2024; Delpuech et al., 2021). Therefore, there is considerable interest in identifying biologically informative intermediate phenotypes that can be reliably measured and integrated into selective breeding programs (Legarra & Christensen, 2023; Goldansaz et al., 2017; Fontanesi, 2016).

Metabolite-profile data, which encompasses the comprehensive profiling of small molecules in biological systems, offers a dynamic snapshot of an organism’s physiological and biochemical state (Fiehn, 2002). Blood metabolite-profile data have shown promise as intermediate phenotypes because they more directly reflect underlying biochemical pathways than traditional production traits (Jang et al., 2019). Since the metabolome displays both genetic and environmental influences, it offers a unique opportunity to bridge the gap between genotypes and complex phenotypes in genomic prediction models (Carmelo et al., 2020). The integration of metabolite-profile data into genetic evaluations has been previously explored in crops such as barley (Guo et al., 2023) and livestock such as pig (Guo et al., 2025). Specifically in the pig application, the study focusing on average daily gain in Duroc pigs has reported small increases in prediction accuracy when including metabolite-profile data collected at the end of the testing period (Guo et al., 2025).

However, the timing of metabolite sampling plays a crucial role (Bromage et al., 2016), and the effects of metabolites on phenotypes may vary across different traits (Aliakbari et al., 2019). Studies such as those by Zhou et al. (2024) and Song et al. (2022) have shown that metabolite-profile data differ significantly by age, breed, and feeding strategy, indicating the value of collecting omics data at multiple physiological stages. Guo et al. (2025) noted that the limited improvement in prediction accuracy might stem from a mismatch between the time of metabolite sampling and the time-integrated nature of average daily gain, highlighting the importance of temporal dynamics in omics-based prediction.

In our study, we address this by analyzing serum metabolite-profile data collected at two developmental stages, 10-week and 20-week of age. This phase span is particularly important, as pigs undergo rapid growth and metabolic transitions during this time, which profoundly influence traits such as feed efficiency and carcass composition. By evaluating five economically important traits, including average daily feed intake (DFI), feed conversion ratio (FCR), backfat thickness (BF), test daily gain (TDG), and loin depth (LD), we provide a more comprehensive assessment of the potential value of metabolite-profile data in pig breeding. We hypothesize that integrating individual-level metabolite-profile data collected at multiple growth stages into genomic prediction models will improve phenotype prediction accuracy compared to using genomic data alone. We further anticipate that the degree of improvement will vary according to the specific trait and timing of metabolite sampling. We also expect that incorporating metabolite-profile data will consistently increase the signals available to genomic prediction models without compromising accuracy.

## Methods

### Dataset

#### Data Collection

Metabolite-profile data was provided by PIC (a Genus company, Hendersonville, TN, USA). Blood samples were taken from 2101 pigs from a single purebred population at both 10-week and 20-week of age and then analyzed using 1H NMR (Nuclear Magnetic Resonance) spectroscopy, providing individual metabolite-profile data. The traits analyzed were: DFI (measured from 10-week to 20-week span), FCR (measured from 10-week to 20-week span), BF (measured at 20-week), TDG (measured from 10-week to 20-week span), and LD (measured at 20-week). Imputed genotypes for a medium density SNP (37k markers) panel were also used for this study.

#### Sampling Strategy

Blood samples were collected from each pig at two distinct timepoints (10-week and 20-week of age) to capture the changes of metabolite-profiles during this time span. This period contains a transition period from nursery to finishing, when environmental features as feed and space change, marking a potential turning point in metabolic activity. Therefore, this stage is particularly relevant for understanding baseline variation in blood serum metabolite-profile data. The finishing phase is also the most feed-intensive period of production, contributing substantially to the total cost of raising pigs. Given the economic and biological importance of this phase, the metabolome at this stage is likely to reflect individual differences in feed efficiency and growth potential. For example, the cost of feeding is very high during the finishing phase, and the individual metabolomic landscape is potentially related to feed efficiency. Therefore, the two sampling timepoints enables capturing both early and late metabolic signatures associated with performance traits.

#### Data Availability

The metabolite-profile data were obtained from blood serum samples. Blood samples were taken and stored under controlled conditions at -80 C to preserve the stability of metabolites prior to analysis. The 1H NMR spectra of the samples were recorded using on a Bruker AV-III 600 MHz spectrometer, which is certified for the In Vitro Diagnostic research (IVDr) analysis, using a BBI probe, and the standardized experiments were done at 310 K. The readings were automatically preprocessed by Bruker software, which performed essential steps such as zero-filling to enhance resolution, apodization to improve signal-to-noise ratio, Fourier transformation to convert time-domain signals to frequency-domain spectra, phase correction and baseline alignment, and spectrum calibration for consistent peak referencing. The comprehensive metabolite-profile data were generated using Bruker IVDr Quantification in Plasma/Serum B.I.Quant-PS™ and Bruker IVDr Lipoprotein Subclass Analysis B.I.LISA™ reports. These advanced quantification platforms enabled automated identification and absolute concentration measurements of a wide range of serum metabolites. The resulting profile included a diverse set of metabolites such as amino acids and their derivatives, ethanol, carboxylic acids, keto acids, sugars, and lipoprotein fractions, among others, capturing a broad overview of each animal’s metabolic state at the time of sampling. The raw metabolites dataset included a total of 149 metabolites, composed of 112 lipoproteins and 37 small-molecule metabolites.

#### Data Quality Control

Quality control (QC) procedures were applied to all datasets used in this study. For metabolite-profile data, metabolites with more than 20% missingness (i.e., measured in less than 80% of the original 2,147 individuals) were excluded. This resulted in a final dataset comprising 78 lipoproteins and 17 small-molecule metabolites, totaling 95 metabolites. For genotype data, standard quality control procedures were applied, including filtering out SNPs with a minor allele frequency (MAF) below 0.01, resulting in 37,199 SNPs retained for analysis. After these QC steps, the final dataset included 1,637 individuals who had complete records for both 10-week and 20-week metabolite-profile data, all five phenotypic traits, and genotype data.

### Model for analysis

#### Heritability Estimation

Heritability of each metabolite was estimated separately for the 10-week and 20-week datasets using the software tool JWAS (Cheng et al., 2018), with each metabolite treated as a quantitative trait. For the five traits (DFI, FCR, BF, TDG, and LD), the proportion of variance explained by each component was also estimated using JWAS (Cheng et al., 2018).

#### Statistical Methods

For the genomic prediction of each trait, we evaluated several linear mixed models that incorporated genotypes and/or metabolite profiles collected at different timepoints, as shown in Table 1. Prior distributions of regression coefficients were specified according to the BayesC method, using two separate sets of hyperparameter values for genotype and metabolite profiles (Wang & Cheng, 2021). When only genotypes were included, the prediction model is expressed as:

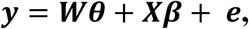

where **y** is the *n* × 1 phenotype vector (n = 1,637 individuals); ***θ*** is the vector of fixed effects and ***W*** is its corresponding design matrix; **X** is the *n* × *p* genotype matrix (p = 37,199 SNPs); **β** is the *p* × 1 vector of marker effects; and ***e*** is the *n* × 1 vector of residual variance for each individual with the distribution of 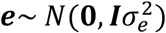.

**Table 1.**
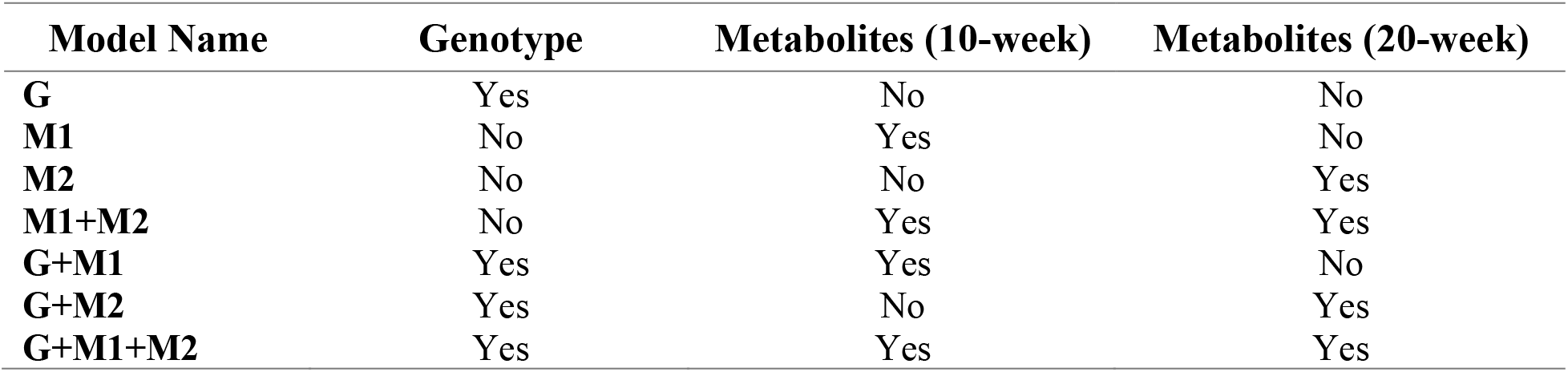
Overview of the seven predictive models constructed using different combinations of genotype and metabolite-profile data. Here, ‘Yes’ indicates inclusion and ‘No’ indicates exclusion.

In the BayesC framework, marker-effect estimation is based on a biologically meaningful assumption that only a subset of all genomic variants has non-zero effects, while the majority do not influence the phenotype. The model therefore infers a null-effect probability, π, which reflects the proportion of SNPs whose effects are zero. For the remaining SNPs, their effect magnitudes are assumed to follow a normal distribution with mean 0 and variance 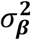 . Thus, the effect of SNP j is modeled as a mixture distribution comprising a point mass at zero and a normal distribution as expressed as:

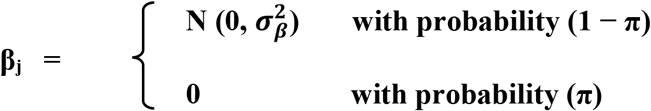

Metabolite-profile data were included as additional random regressors. When metabolites from 10-week and 20-week were analyzed together, the model became:

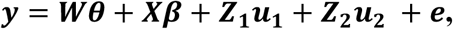

where variables ***y, Wθ, Xβ***, and ***e*** are the same as those in the conventional BayesC model as described above, ***Z***_**1**_ and ***Z***_**2**_ are the matrices of metabolites from weeks 10 and 20, respectively, where each matrix entry in the *i*^th^ row and *j*^th^ column represents the *j*^th^ metabolite feature for individual *i*, and ***u***_**1**_ and ***u***_**2**_ are their corresponding effect vectors. For DFI and TDG, litter was included as random effect and birth year-month was included as fixed effect, whereas for FCR, BF, and LD, only litter was included as random effect. The choice of including random and fixed effect in the models for each trait was based on their optimal prediction performance in the baseline genotype-only (G) model.

Gibbs samplers were applied to obtain samples of the joint posterior distributions of components within each model. Samples were saved every 5^th^ iteration, yielding 1,000 posterior samples after 5,000 total iterations. Convergence was verified by visual inspection of trace plots for key parameters.

#### Validation Method

Prediction accuracy was evaluated as the Pearson correlation between the estimated values and adjusted phenotypes in a testing set. Model performance was compared across the seven models. To evaluate prediction accuracy, the full dataset was randomly divided into a training set (80%) and a testing set (20%). We ensured that all levels of litter and birth year-month were observed in both training and testing sets. Model parameters were estimated using the training set and estimated values were calculated for individuals in the testing set. This procedure was repeated for 20 replicates.

The Figure 1 illustrates the whole process including metabolite-profile data collection procedure and implementation of these datasets in the genomic prediction models.

**Figure 1.**
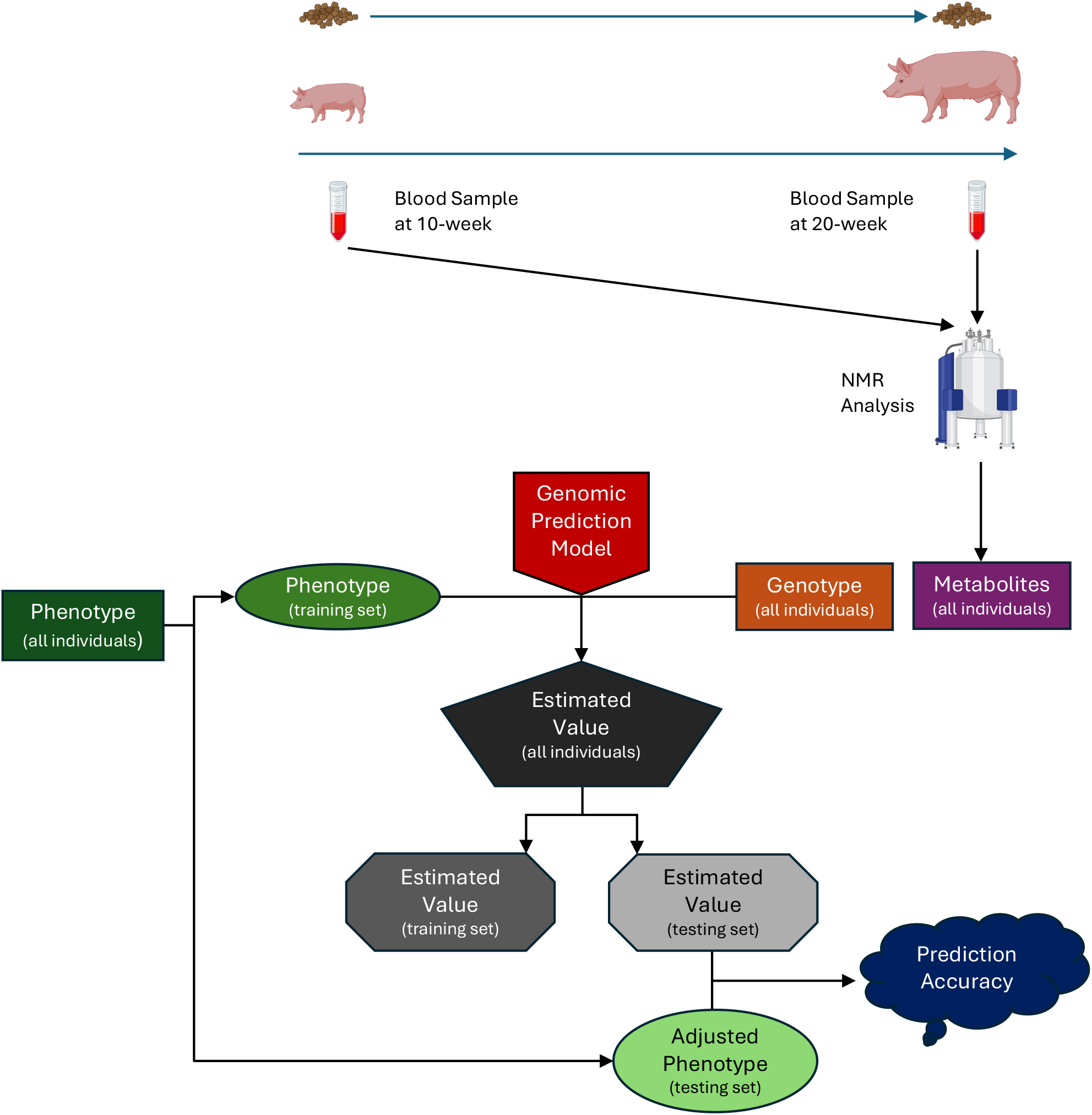
A detailed workflow from the data collection to the prediction accuracy estimation.

## Results

### Heritability Estimation of Metabolites

To assess the genetic basis of serum metabolites variation, we estimated the heritability (h^2^) of 95 metabolites at 10-week and 20-week. The goal of this analysis was to evaluate whether these metabolites are heritable, and to quantify their heritability.

### Distribution of heritability at each timepoint

Figure 2 (a) illustrates the distribution of heritability (h^2^) estimates for serum metabolites collected at 10 and 20 weeks of age. It shows that the most frequent values for the 10-week heritability (reddish-bars) are mostly within 0.07 to 0.30, which is relatively higher than the 20-week heritability (greenish-bar), which shows values concentrated between 0.04 and 0.28. This pattern suggests that, as pigs progress through the finishing phase, their metabolite profiles exhibit decreasing relative genetic influence and increasing environmental contributions.

**Figure 2.**
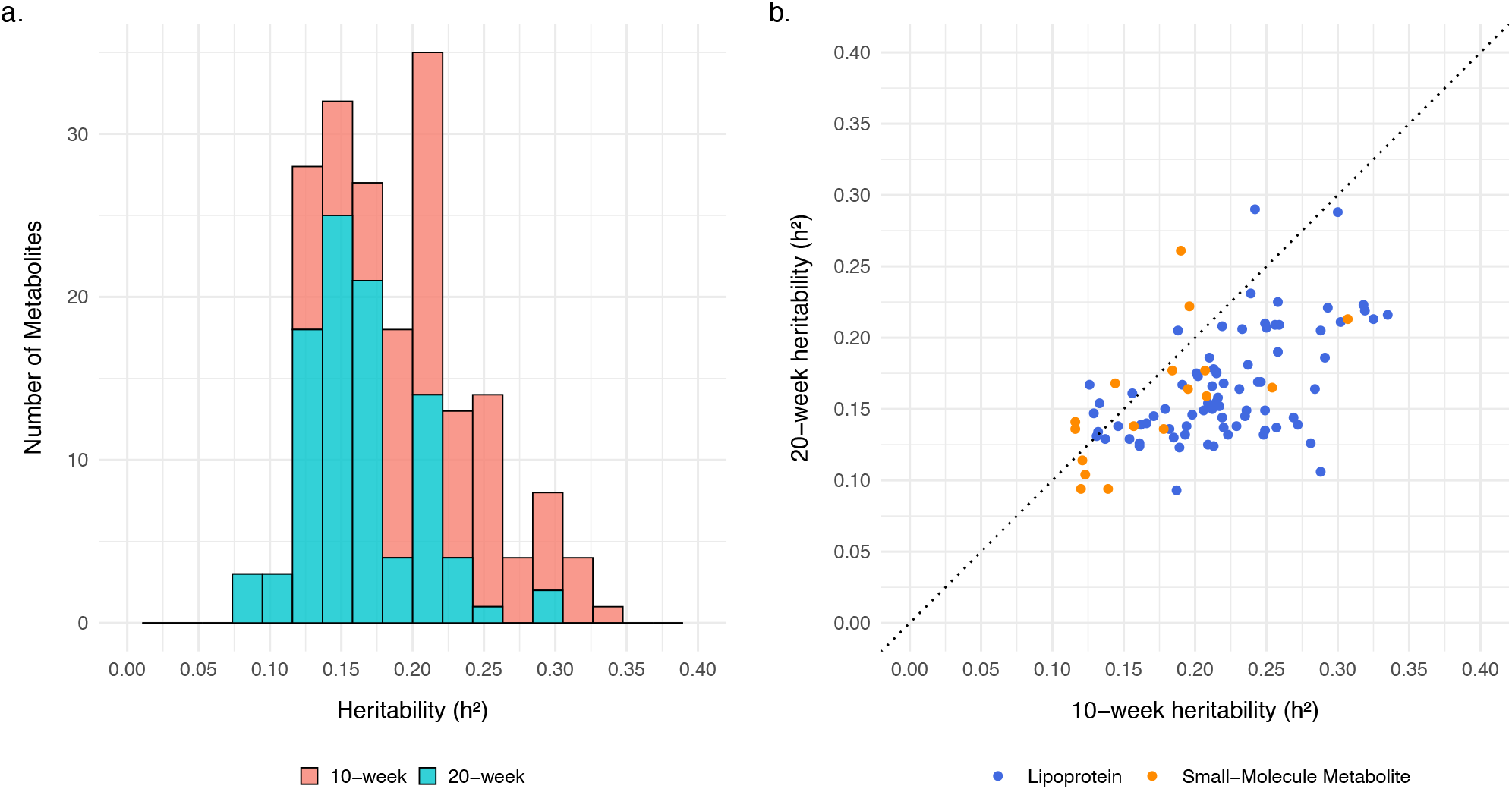
Estimated heritability of metabolites. **a**. Distribution of heritability estimates for metabolites measured at 10-week and 20-week. **b**. Scatter plot comparing heritability estimates for individual metabolites between the 10-week and 20-week sampling points. Points above the diagonal line indicate higher heritability at 20-week, whereas points below suggest higher heritability at 10-week.

Figure 2 (b) compares the heritability values of individual metabolites across the two stages. Points above the diagonal line indicate higher heritability at 20 weeks, whereas points below suggest higher heritability at 10 weeks. Most metabolites lie close to or below the diagonal, suggesting that genetic influence generally decreases or remains stable with age. Notably, lipoproteins (blue dots) are more frequently positioned below the diagonal line compared to small molecule metabolites (orange dots), indicating that lipoproteins, in particular, may be more consistently heritable at 10-week.

### Proportion of variance explained by genotype and metabolites component for five traits

Table 2 shows that the proportion of variance explained increased when genotype and metabolite-profile data were combined. For FCR and BF, the highest performance was observed with the G+M2 model, and for DFI, TDG, and LD, the best performance was achieved with the G+M1+M2 model.

**Table 2.**
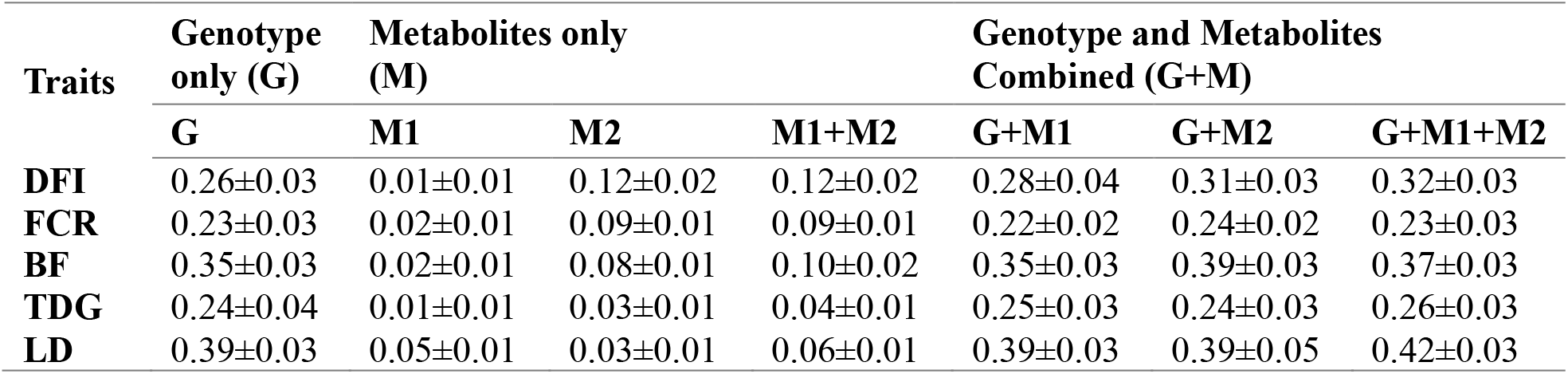
Proportion of variance explained by genotype and metabolite components for five traits across different models (h^2^±SD)

| Traits | Genotype only (G) | Metabolites only (M) |  |  | Genotype and Metabolites Combined (G+M) |  |  |
| --- | --- | --- | --- | --- | --- | --- | --- |
|  | G | M1 | M2 | M1+M2 | G+M1 | G+M2 | G+M1+M2 |
| **DFI** | 0.26±0.03 | 0.01±0.01 | 0.12±0.02 | 0.12±0.02 | 0.28±0.04 | 0.31±0.03 | 0.32±0.03 |
| **FCR** | 0.23±0.03 | 0.02±0.01 | 0.09±0.01 | 0.09±0.01 | 0.22±0.02 | 0.24±0.02 | 0.23±0.03 |
| **BF** | 0.35±0.03 | 0.02±0.01 | 0.08±0.01 | 0.10±0.02 | 0.35±0.03 | 0.39±0.03 | 0.37±0.03 |
| **TDG** | 0.24±0.04 | 0.01±0.01 | 0.03±0.01 | 0.04±0.01 | 0.25±0.03 | 0.24±0.03 | 0.26±0.03 |
| **LD** | 0.39±0.03 | 0.05±0.01 | 0.03±0.01 | 0.06±0.01 | 0.39±0.03 | 0.39±0.05 | 0.42±0.03 |

### Prediction Accuracy Estimation

Table 3 presents a comparison of prediction accuracy for five economically important traits across multiple predictive models. The genotype-only (G) model provided a moderate-to-high baseline prediction accuracy. Prediction accuracies using metabolites-only models (M1 and M2) were consistently lower than the G model for all traits, except for DFI and FCR, where M2 matches G for FCR and surpasses it for DFI. Among these metabolite-only models, the model with 20-week data (M2) yielded higher accuracies than the model with 10-week data (M1) for DFI, FCR, BF, and TDG. In contrast, the reverse pattern was observed for LD, although the standard-error intervals for TDG and LD overlap. Combining both timepoints (M1+M2) clearly improved prediction over M1 with overlapping SEs for TDG and LD. In contrast, prediction with M1+M2 improved slightly over M2 for BF and LD, with overlapping SEs, and showed no improvement for DFI, FCR or TDG. However, except for DFI and FCR, the resulting accuracies with M1+M2 remained inferior to the G model. It is worth noticing that for DFI, the M1+M2 model achieved a higher prediction accuracy than G.

**Table 3.** Comparison of prediction accuracy for five traits across genomic and metabolomic prediction models (Mean±SD).

| Traits | Genotype only (G) | Metabolites only (M) |  |  | Genotype and Metabolites Combined (G+M) |  |  |
| --- | --- | --- | --- | --- | --- | --- | --- |
|  | G | M1 | M2 | M1+M2 | G+M1 | G+M2 | G+M1+M2 |
| **DFI** | 0.31±0.04 | 0.09±0.06 | 0.32±0.05 | 0.32±0.05 | 0.31±0.04 | 0.41±0.04 | 0.41±0.04 |
| **FCR** | 0.27±0.05 | 0.13±0.05 | 0.27±0.05 | 0.27±0.05 | 0.28±0.05 | 0.33±0.06 | 0.33±0.06 |
| **BF** | 0.41±0.04 | 0.11±0.03 | 0.23±0.05 | 0.25±0.05 | 0.41±0.04 | 0.45±0.04 | 0.45±0.04 |
| **TDG** | 0.28±0.05 | 0.08±0.05 | 0.16±0.04 | 0.16±0.05 | 0.28±0.04 | 0.32±0.04 | 0.32±0.04 |
| **LD** | 0.42±0.04 | 0.17±0.06 | 0.15±0.05 | 0.19±0.05 | 0.44±0.04 | 0.44±0.04 | 0.45±0.04 |

Integrating metabolite-profile data with genotype data (G+M) mostly improves prediction accuracy compared to using genotype (G) alone, although the magnitude of improvement varies by trait and metabolite sampling timepoint. For instance, an increase in prediction accuracy over the G model is observed for all five traits when using the G+M2 and G+M1+M2 models, whereas the G+M1 model does not yield similar improvements. Compared to all metabolites-only models, combining any metabolite data and genotype information into the same model leads to equal or improved performance in all cases except for DFI, where M2 and M1+M2 provided higher accuracy than G+M1. It is also worth noticing that the G+M1+M2 model distinctively achieved the highest prediction accuracy among all seven models only for LD, whereas for DFI, FCR, BF, and TDG, G+M1+M2 gives similar performance to G+M2.

Figure 3 shows the Monte Carlo cross-validation distribution of predictive accuracies for all models relative to G used as baseline. Each point represents, for one of 20 random train/test splits, the differences in prediction accuracy between the model indicated on the x-axis and the genotype-only (G) baseline. In these box plots, two distinct colors represent different model categories: blue indicates metabolites-only models (i.e., M1, M2, and M1+M2), and yellow indicates models combining genotypes and metabolites (i.e., G+M1, G+M2, G+M1+M2). The box plots represent the distributions of accuracy differences relative to G for all models and traits. These plots provide additional insight into how metabolite-profile data, either alone or in combination with genotype, impacts prediction accuracy relative to the genotype-only baseline.

**Figure 3.**
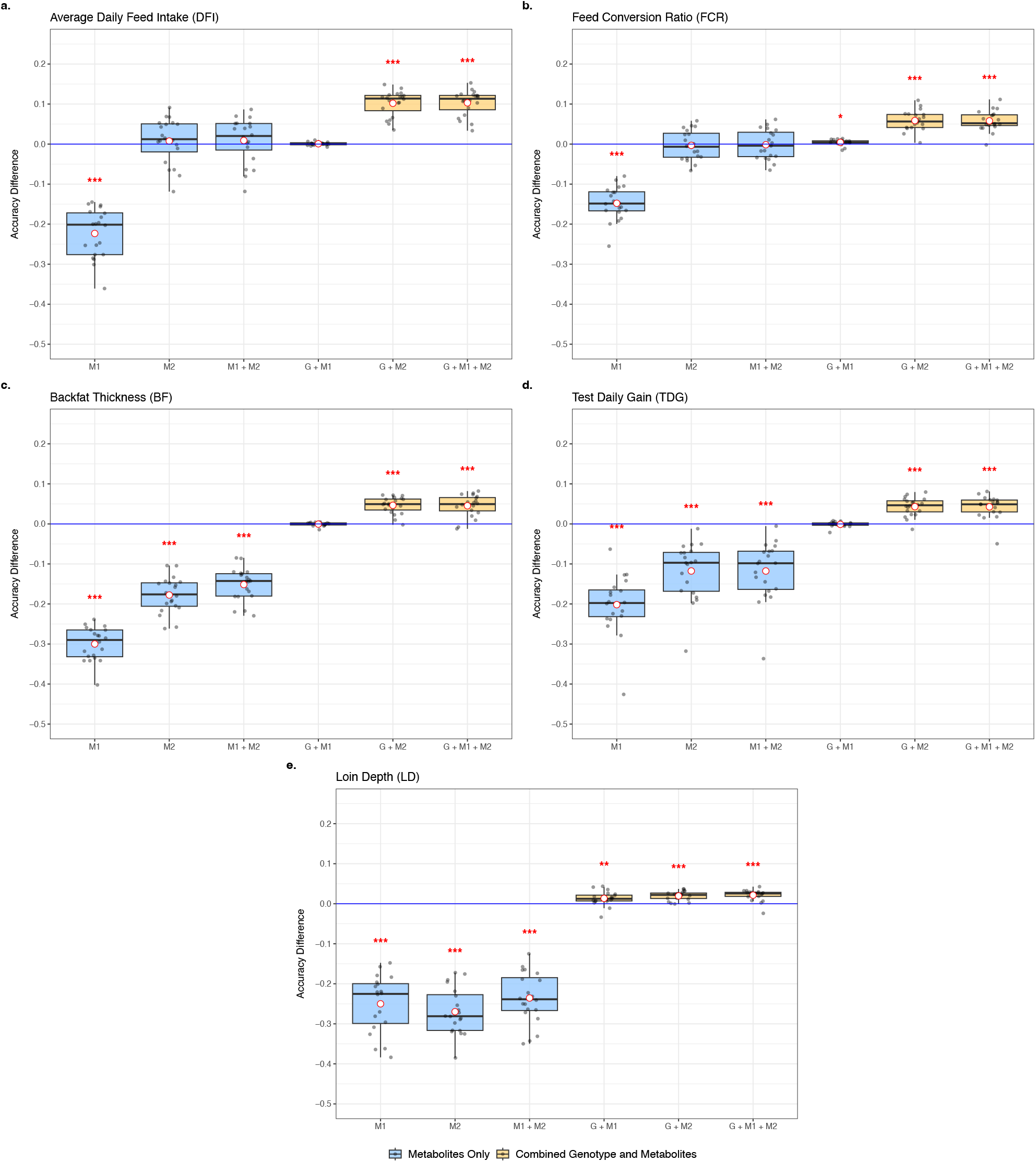
Box plots showing the differences in prediction accuracy across various models relative to the genotype-only baseline (Model G) for five traits: **a**. DFI, **b**. FCR, **c**. BF, **d**. TDG, and **e**. LD. Each plot represents the difference in prediction accuracy between the genotype-only model and metabolites-only models (blue boxes) or combined genotype and metabolites models (yellow boxes). Statistical significance is indicated by * (*p-value < 0.05, ** p-value < 0.01, *** p-value < 0.001).

All metabolites-only models (M1, M2, and M1+M2) showed notably lower (p<0.001) prediction accuracy across traits compared to the genotype-only baseline, except for traits DFI and FCR, where the prediction accuracy using only the 20-week (M2) and the combined 10-week and 20-week (M1+M2) metabolite-profile data were similar (p>0.05) to that of the G model. This suggests that late-stage metabolite-profile data alone may be sufficient to provide an alternative predictor for these two traits, with performance similar to using genomic data, when the latter data is not available. This finding highlights the potential utility of metabolite-profile data (especially late-stage) as indicators for specific economically important traits.

Furthermore, M2 showed a clear superiority as a single timepoint across traits than M1, except for LD. Also, combining metabolite-profile data from both M1 and M2 mostly yielded higher prediction accuracies than either timepoint alone with some exceptions. This suggests that metabolite-profile data from different developmental stages capture complementary biological information, thus providing an enhanced predictive ability.

Integrating metabolite-profile data with genotype information generally enhanced prediction accuracy, though the magnitude of the improvements varied depending on the trait and the age of metabolite sampling. Specifically, the inclusion of 20-week metabolite-profile data G+M2 consistently increased accuracy (p<0.001) across all five traits, highlighting the importance of the late-stage metabolite-profile data. Incorporating only the 10-week metabolite-profile data (G+M1) did not improve prediction accuracy (p>0.05) over the genotype-only model for DFI, BF, and TDG, whereas it provided noticeable gains for FCR (p<0.05) and LD (p<0.01). Combining metabolite-profile data from both timepoints (G+M1+M2) achieved a significant overall improvement in accuracy (p<0.001) relative to the G model across all five traits. This indicates that including both early and late metabolite-profile data in the model allows it to explore complementary biological signals that are beneficial for the prediction of phenotypes.

The influence of each combination of type of information included in the model on the predictive performance may reflect how each trait has a distinct relationship dynamic with the blood metabolomic landscape. Notably, for DFI, both M2 and M1+M2 outperform G, indicating that late-stage metabolite-profile data may serve as an alternative predictor. For FCR and LD, combining metabolite-profile data from either timepoint with genotype data improved prediction accuracy with the highest gain achieved while using G+M2 and G+M1+M2. For LD, the greatest gain observed when using both timepoints together with genotype data. This finding implies that these two traits might be a cumulative trait benefiting from metabolite information across multiple developmental stages. On the other hand, for BF and TDG, prediction accuracy differences between G and G+M1 were statistically non-significant (p>0.05). However, for these two traits, both G+M2 and G+M1+M2 models provided the highest prediction accuracy with a significance level of p<0.001. All these patterns suggests that late-stage metabolite-profile data captured biologically more relevant variation for predicting these five traits.

## Discussion

Our study evaluated the potential of integrating individual-level metabolite-profile data collected at multiple growth stages from blood serum into genomic prediction models for five economically important traits in pigs: DFI, FCR, BF, TDG, and LD. Our findings confirm that many metabolites are moderately heritable and combining them with genotype data led to consistent improvements in prediction accuracy across all traits. These findings align with previous studies. For example, Aliakbari et al., (2019) showed that certain metabolites are moderately to highly heritable and genetically correlated with growth traits such as body weight and average daily gain, suggesting their value as intermediate phenotypes. In addition, our findings emphasize that the benefit of incorporating metabolite-profile data across different developmental stages into genomic prediction models must be evaluated on a trait-by-trait basis. Importantly, integrating metabolite-profile data consistently complemented genomic predictions without negatively affecting accuracy across all traits.

The biological relevance of incorporating metabolite-profile data in genomic prediction is trait dependent. For traits like BF and TDG, the inclusion of late metabolite-profile data (20-week) provides further useful signals when combined with genotype data. DFI, FCR and LD, being influenced by both early and late biological stages, benefit from the integration of metabolite-profile data from multiple timepoints. Furthermore, for LD, the highest performance was achieved distinctively while combining both timepoint metabolite-profile data with genotypes. These results are consistent with previous research showing that metabolite profiles are influenced by breed, nutritional environment, and physiological stage (Liu et al., 2021), which underscores the importance of aligning sampling strategy with traits. These findings support the targeted use of timepoint-specific metabolite-profile data for trait-informed breeding strategies.

Heritability estimates for the metabolites ranged from low to moderate at both 10-week and 20-week timepoints. A similar range was reported by Aliakbari et al. (2019) for serum metabolites in dairy cattle, supporting the hypothesis that metabolite levels are genetically influenced. Our analysis showed that heritability estimates were slightly higher at 10-week, and the metabolite-profile data from 20-week were more useful in improving predictive performance when combined with genotypes for most traits. This aligns with observations from Rohart et al. (2012), who reported that metabolite-profile data reflect distinct metabolic states and may be more informative for predictive performance at certain physiological stages. In addition, genome-wide association studies and gene–metabolite network analyses have identified candidate biomarkers and biological pathways related to feed efficiency, further supporting the potential role of metabolite-profile data in improving trait prediction and genetic evaluation related to feed efficiency (Wang and Kadarmideen, 2020).

In the current analysis, we focused on phenotype prediction. Our results indicate that incorporating early-stage metabolite-profile data (10-week), which exhibit higher heritability, may effectively capture biologically meaningful genetic variation that is useful in genomic prediction models. Nonetheless, higher prediction accuracies were generally observed when incorporating late-stage (20-week) metabolite data. This pattern likely reflects the proximity of late-stage metabolite sampling to the actual phenotypic measurements, providing a more immediate snapshot of physiological status. In future studies, we plan to evaluate the utility of metabolite-profile data specifically for predicting breeding values, directly informing genetic improvement in livestock populations.

## Conclusions

This study demonstrates that integrating individual-level metabolite-profile data with genotype data can improve the prediction accuracy for average daily feed intake, feed conversion ratio, backfat thickness, test daily gain, and loin depth, which are key production traits in pigs. While metabolite-profile data alone yielded relatively low predictive power in most scenarios, likely due to the complex and temporally dynamic nature of the metabolome, their combination with genotype data consistently enhanced prediction performance across all traits. The moderate heritability observed for many metabolites further supports their potential as intermediate phenotypes that reflect both genetic and physiological processes. By integrating this omics layer, prediction models gained access to non-redundant biological information not captured by genotypes alone. Future studies should focus on optimizing the timing and selection of metabolites, leveraging trait-specific biological knowledge, and integrating metabolomic information into large-scale genomic evaluation programs. Specifically, scenarios in which metabolite data are available only for a subset of animals should be explored using sophisticated modeling approaches, such as NNMM (Zhao et al., 2022), to effectively incorporate partial metabolomic information.

## List of abbreviations

DFI: Average Daily Feed Intake
FCR: Feed Conversion Ratio
BF: Backfat Thickness
TDG: Test Daily Gain
LD: Loin Depth
G: Genotype-only model
M1: Metabolite-only model when metabolite-profile data is from 10-week
M2: Metabolite-only model when metabolite-profile data is from 20-week
M1+M2: Metabolite-only model when metabolite-profile data is from both 10-week and 20-week
G+M1: Combined Genotype and Metabolite model when metabolite-profile data is from 10-week
G+M2: Combined Genotype and Metabolite model when metabolite-profile data is from 20-week
G+M1+M2: Combined Genotype and Metabolite models when metabolite-profile data is from both 10-week and 20-week

## Declarations

### Ethics approval and consent to participate

Not applicable

### Consent for publication

Not applicable

### Availability of data and materials

The data explored in this study is available upon request to PIC

### Competing interests

The authors declare that they have no competing interests.

### Funding

This work was supported by USDA-NIFA awards 2023-67015-39564, and 2023-70412-41054. Instrumentation was supported by a grant from NIH (S10 RR025677) for the console upgrade, a Vanderbilt Trans-Institutional Programs (TIPs) grant to purchase the IVDr equipment, NIH supplement (3R35GM118089-04S1) for the helium liquefier, all accompanied by Vanderbilt University matching funds.

### Authors’ contributions

HC conceived the study. QAHR implemented and performed the statistical analyses with the assistance or guidance from HC, JQ, DL (Dingzhen), BV, CYC, JH, ERC, and DL (Daniela). HC and JQ contributed to the experimental design. QAHR, JQ, and DL (Dingzhen) drafted the manuscript. All authors read and approved the final manuscript.

## Acknowledgements

Not Applicable

## Additional Files

**Additional file 1 Figure S1** A comparison of prediction accuracy distribution across models and traits.

**Additional file 1 Figure S1:**
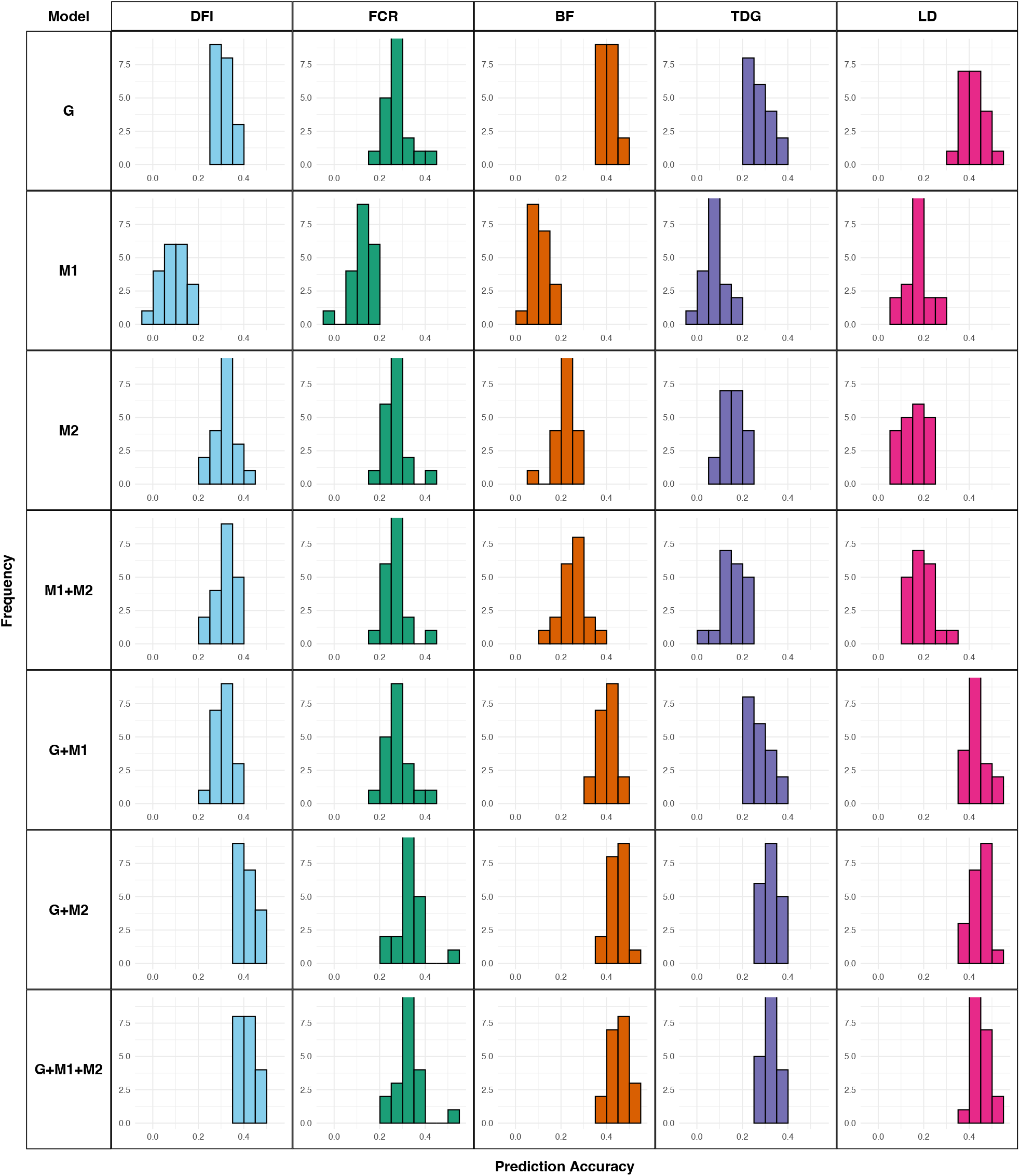
Distribution of prediction accuracy for five traits — FIP, FCR, BF, TDG, and LD — across seven genomic and metabolomic prediction models. The models include genotype-only (G), 10-week metabolite-only (M1), 20-week metabolite-only (M2), combined 10- and 20-week metabolites (M1+M2), genotype + 10-week metabolites (G+M1), genotype + 20-week metabolites (G+M2), and genotype + both 10- and 20-week metabolites (G+M1+M2). Each histogram shows the frequency distribution of prediction accuracy values obtained from cross-validation replicates (20 replications each) for the respective trait and model combination. Different colors represent different traits such as sky-blue for FIP, green for FCR, orange for BF, violet for TDG, and pink for LD.

## Notes

### Competing Interest Statement

The authors have declared no competing interest.

